# Transcriptional plasticity of fast core chromosomes governs establishment of a fungal pathogen on evolutionarily distant plant lineages

**DOI:** 10.1101/2025.03.27.645276

**Authors:** Vidha Srivastava, Siddharth Kaushik L S, Naga Jyothi Pullagurla, Bernd Zechmann, Antonio Di Pietro, Debabrata Laha, Amey Redkar

## Abstract

The Fusarium oxysporum (Fo) species complex encompasses a diverse range of filamentous plant pathogens, some of which provoke systemic infections in angiosperms, leading to vascular wilt disease. Current understanding of the Fo pathogenicity mechanisms is primarily centered on vascular plants, where individual isolates of Fo exhibit contrasting lifestyles across the endophyte-pathogen continuum on different host species. Although, Fo isolates were recently shown to cause disease on the non-vascular liverwort Marchantia polymorpha (Mp), the transcriptional control of Fo effectors during infection of this bryophytic host is largely unknown. Here, we took a comparative transcriptomic approach to ask how different Fo isolates with contrasting interaction outcomes on angiosperms (pathogenic versus endophytic) adapt to a distantly related plant lineage lacking xylem. We found that the core effector complement encoded on genomic regions shared across all Fo isolates are actively transcribed in Mp, whereas effectors encoded on lineage-specific (LS) genomic regions that contribute towards host-specific pathogenicity in angiosperms are not. Moreover, we observed enhanced transcriptional activation of effector clusters located on the three highly syntenic fast core chromosomes during growth on Mp, as well as divergent lineages and lifestyles, suggesting a conserved role in plant associations. Loss of a compatibility-associated effector encoded in a core effector cluster led to misregulation of other effectors in core clusters. Our findings reveal an unexpected role of fast core chromosomes in determining compatibility of Fo across a broad spectrum of plant lineages and establishes evolutionarily conserved gene networks essential for fungus-plant associations.

## INTRODUCTION

Microbial associations have played a crucial role in the evolutionary history of plants and their adaptation to early terrestrial ecosystems [1]. Paleobotanical evidence from the Rhynie chert revealed ancient fungal-plant associations, observed as fungal fruiting bodies likely from ascomycetous fungi, in the sub-epidermal layers of the primitive land plant *Asteroxylon* [2]. Moreover, early Devonian plant fossils also showed associations with fungus-like organisms likely depicting cell-wall associated responses during the relationship of extant plants with intruding filamentous microbes [3,4]. Specialized anatomical features such as a vascular system for conducting water and minerals, differentiate tracheophytes (vascular land plants) from evolutionarily distant non-vascular plant lineages. This innovation has since led to the accelerated colonization of diverse habitats by plants, contributing to their evolutionary success. Moreover, such adaptations have also driven complex interactions with microorganisms leading to parasitic or mutualistic relationships [5]. While the plant vasculature has primarily contributed to mechanical support and long-distance transport, it has also been exploited by co-evolving microbes for systemic infections and long-distance spread across their hosts [6].

A particularly destructive group of filamentous microbes adapted to cause systemic infections are vascular wilts, which colonize the inner, nutrient poor zone of the xylem. Among these, *Fusarium oxysporum* (Fo) constitutes a species complex with a broad host range of tracheophytes that causes devastating losses in over 150 crops [7,8]. Molecular clock estimates in geological timescale pinpoint the origin of the genus *Fusarium* around 91.3 million years ago (mya) coinciding with the radiation of angiosperms [9] whose further divergence contributed to host specificity about 65 mya ago in the cretaceous period [10].

The Fo species complex (FOSC) provokes systemic infections through the xylem causing wilt disease [7,11]. Adaptation to the xylem niche is determined by accessory or lineage-specific (LS) genomic regions which confer pathogenicity and wilting [12,13]. The accessory chromosome 14 of the tomato pathogenic isolate Fo f.sp. *lycopersici* (Fol) encodes host-specific virulence proteins, initially discovered from infected xylem sap and are termed as <u>S</u>ecreted In <u>X</u>ylem (SIX) effectors [14]. The entire chromosome 14 of Fol can be transferred across isolates to gain pathogenicity on tomato confirming its role in host-specific adaptation [12]. Despite exhibiting host-specific pathogenicity, most isolates of Fo can colonize roots of diverse plants asymptomatically as endophytes [15]. Different lifestyles of Fo result from the activity of virulence factors and are associated with major transcriptional shifts between core and LS effector complements coordinating establishment of the fungus in the root cortex or transition to the xylem [15].

Our current understanding of vascular wilt pathogens and host responses comes primarily from flowering plants. However, a previous study on fungal endophytes of the non-vascular plant *Marchantia polymorpha* (Mp) showed the presence of Sordariomycetes including Fo [16]. Research on non-vascular plant models provides opportunities to understand evolution of molecular plant-microbe interactions (Evo-MPMI) and plant immunity [17–21]. Such comparative studies across fungal and plant lineages provide insights into the emergence of systemic infections and niche-specific adaptation. Establishment of a Fo-Mp pathosystem revealed conserved infection strategies independent of host-specific mechanisms in the fungus during colonization of a non-vascular host [21]. This includes the conserved role of <u>E</u>arly <u>R</u>oot <u>C</u>olonization (ERC) effectors deployed on evolutionarily distant land plants and the specific role of SIX effectors only in tracheophytes [15,21]. Together, these findings pointed towards differences in transcriptional regulation of virulence factors encoded on core and LS regions during infection of Fo on distinct hosts.

Compartmentalized two-speed genomes are a hallmark of fungal phytopathogens that drives their adaptive co-evolution with the plant host [15,22]. An open question is how these distinct genomic regions coordinate the infection process and whether such coordination contributes to host range expansion or adaptation to the tissue-specific niches. Here we used comparative transcriptomics during Fo infection on vascular and non-vascular hosts to study host-specific regulation of core and LS effector genes. We find that fast core chromosomes regulate plant associations across vascular and non-vascular lineages as well as in different Fo isolates. Transcriptional activation of a set of five effector gene clusters on fast core chromosomes are guiding with fungal associations across evolutionarily divergent hosts and lifestyles. Loss of a core effector impaired the infection program by misregulating the effector gene clusters on fast core chromosomes. Our results reveal an unexpected role of fast core chromosomes in establishing compatibility across divergent land plant lineages.

## RESULTS

### Fo shows a comparable development during vascular host and bryophytic host *M. polymorpha*

Previous studies had shown that Mp thalli inoculated with different Fo isolates show macroscopic disease symptoms including chlorosis and maceration with a localized infection at the centre of the thalloid body [15,21]. To determine the dynamics of Fo development, Mp accession Tak-1 was inoculated using a modification of a previously established protocol [21] (see materials and methods). At 3 days post inoculation (dpi), Mp thalli inoculated with the tomato pathogenic isolate Fol or the endophytic isolate Fo47, showed minor disease symptoms, primarily chlorosis (Figure. 1a). Quantification of the fungal biomass in the infected thalli at 3 dpi did not show detectable differences between different Fo isolates (Supplemental Figure. 1). The first signs of plant cell death became visible at 4 dpi resulting in browning of the thalli. Maceration of the thallus tissue was observed at 5 dpi compared to mock, indicating progressive cell death (Figure. 1a, b; one-way Avova Tukeýs Post hoc HSD test, p-value = 0.0001).

**Figure. 1.**
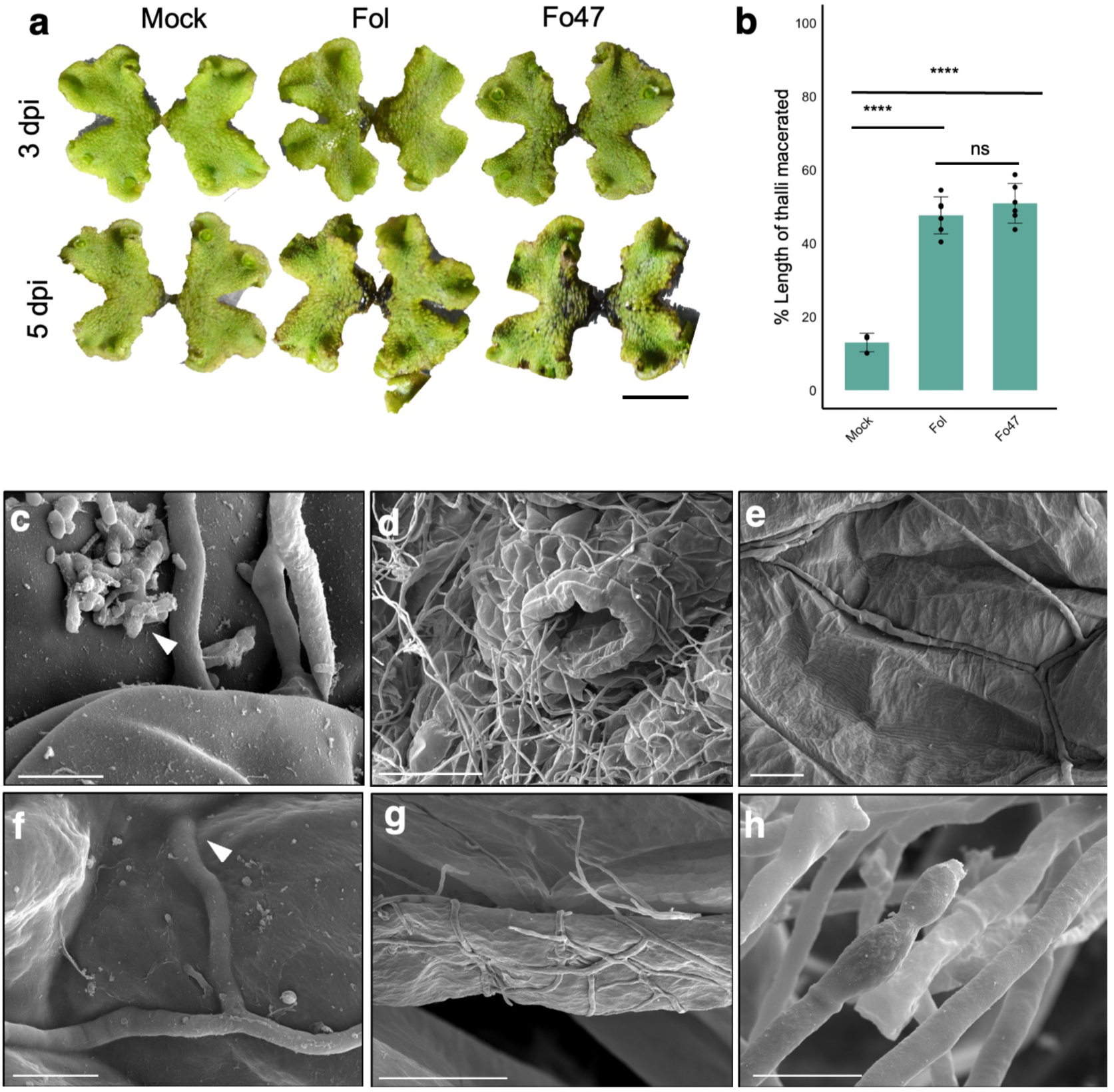
Different isolates of *F. oxysporum* infect *M. polymorpha* and complete their infection cycle. **a,** Macroscopic images showing disease symptoms on *M*. *polymorpha* Tak-1 male plants after dip inoculation of the agar block with 10^5^ microconidia ml^−1^ of the *Fusarium oxysporum* (Fo) strains Fol (tomato pathogen) or Fo47 (endophyte), or water (mock). Images taken at the indicated time points (days post-inoculation, dpi) represent three independent experiments. Bar, 1 cm. **b,** Bar plot representing the per cent length of thallus macerated as measured from the centre to the apical notch (n ≥3 thalli). Each dot represents a measurement from an individual thallus and error bars represent the standard deviation from the mean. Statistical significance was calculated by performing one way ANOVA followed by Tukey’s *post hoc* HSD test. ns = non significant (p-vaue > 0.05) **c-h,** and Scanning Electron Microscopy (SEM) shows different stages of the infection cycle of *F. oxysporum on* Tak-1 at 3 dpi. **c,** germinated microconidia; **d,** colonisation of the thallus through air pore; **e,** intercellular colonization on the thallus surface; **f,** direct penetration through the cuticle (white arrowhead); **g,** hyphae entangled on the rhizoid without showing any intracellular growth; **h,** Phialide-like structures formed on the surface of the colonized thalli. (c, f, h: scale bar = 5μm; d: bar = 50μm; e: bar = 10μm; g: bar = 40μm).

To further characterise the interaction between Mp Tak-1 and Fo we performed a Scanning Electronic Microscopy (SEM) of infected samples at 3 dpi, which captured all the representative infection modes of microconidia germination (Figure. 1c), fungal colonization and establishment, including entry via air pores (Figure. 1d), penetration through the cell-cell junctions (Figure. 1e), or direct penetration by rupturing the epidermal cell (Figure. 1f). Fungal colonization was also visible on the surface of rhizoids, although development of the mycelium inside the rhizoids was rarely observed (Figure. 1g). By 3 dpi the thallus surface was covered with mycelium (Figure. 1d) and SEM analysis also revealed the fungal fruiting body-like structures (Figure. 1h) indicating the completion of the Fo lifecycle on Mp. These results indicate that the 3 dpi timepoint provides a representative stage of infection for monitoring transcriptional responses of the host and the pathogen.

### Core genomic regions of Fo mediate land plant associations while LS regions determine access to the xylem niche

Previous work has shown that core pathogenicity mechanisms are likely sufficient for Fo establishment on Mp whereas, host-specific virulence determinants are largely dispensable for infection on this non-vascular host [21]. To understand the unknown transcriptional network that contributes to establishing fungal association on non-vascular and vascular hosts, we performed RNA-seq analysis of 21-day-old Mp Tak-1 thalli infected with Fol at 3 dpi and compared it with the previously generated transcriptome of tomato roots infected with Fol at 3 dpi as well as with the axenically grown Fol culture [15].

Differential expression analysis with log2FC > 2 or < -2 along with padj < 0.05 of axenically grown Fol cultures versus infected Mp thalli releaved a transcriptional shift of a total of 4,721 genes, which is similar to the previous dataset of the Fol transcriptome from infected tomato roots which detected 3,179 differentially expressed genes (DEGs) versus axenically grown cultures. (Figure. 2 a,b and Supplemental Figure. 2a). Around 32% of the upregulated DEGs of Fo were shared between infection of the non-vascular Mp and the vascular host tomato (Supplemental Figure. 2a,b). Furthermore, around 26% of downregulated Fo DEGs were shared on both hosts (Supplemental Figure. 2a,b). Gene ontology (GO) enrichment analysis of the common DEGs using an FDR of 0.05, showed an enrichment of enzymatic functions such as pectate lyase, polysaccharide degradation, glycosidase, hydrolase and membrane transporters in the positively regulated geneset (Supplemental Figure. 2c). Moreover, thiamine, ATP and nucleoside metabolism, were shared downregulated processes (Supplemental Figure. 2d). While a significant subset of downregulated genes shared between Mp and tomato, 58% of the genes upregulated during Mp colonization were unique, suggesting the involvement of host-specific fungal responses operating alongside a core arsenal for wilting on angiosperms (Supplemental Figure. 2b).

**Figure. 2.**
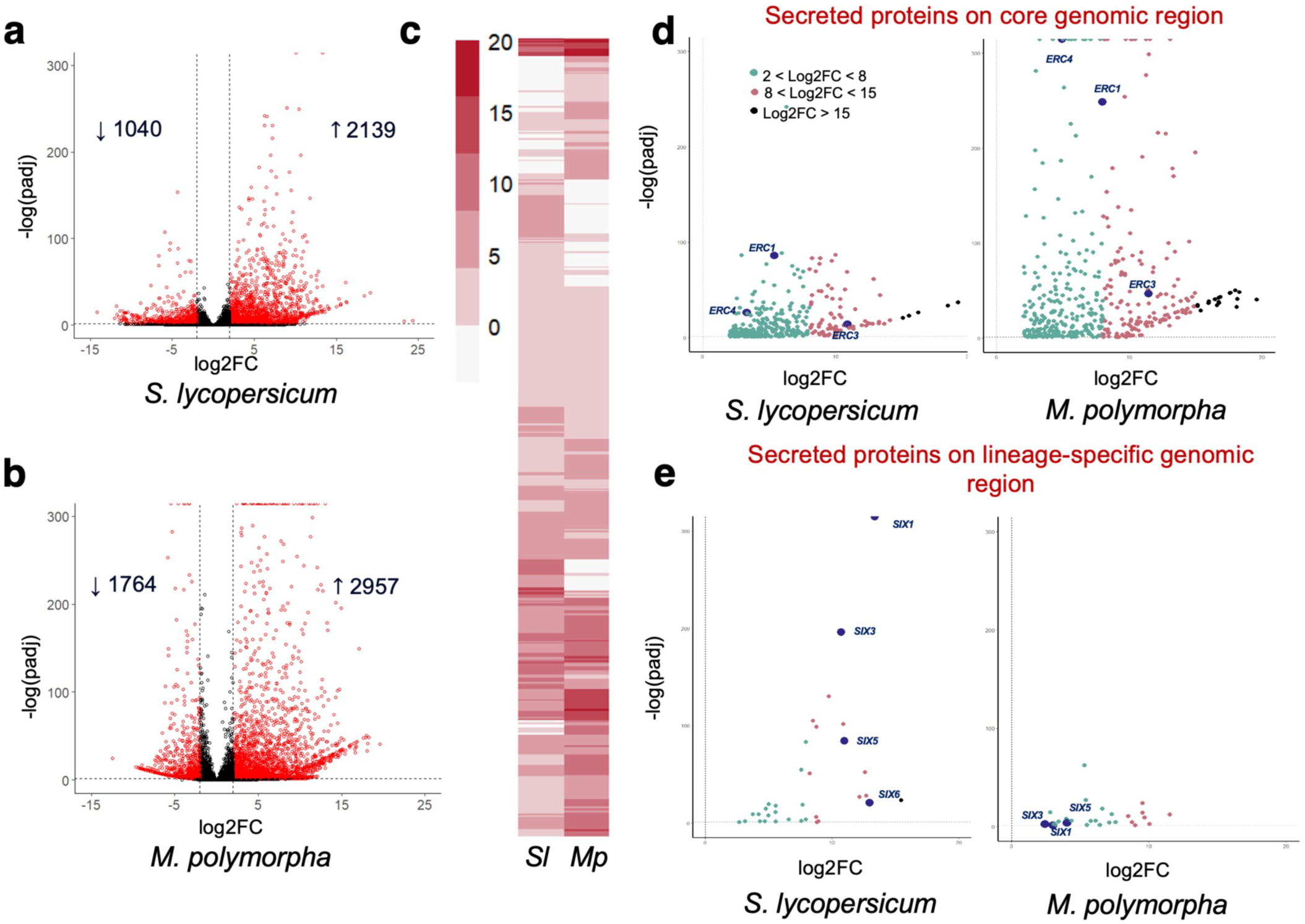
Transcriptional regulation of core and lineage-specific genomic regions of *F. oxysporum* during infection of evolutionarily distant plant lineages. a-b, Volcano plot showing pairwise differential expression analysis of Fol genes at 3 dpi during growth on tomato roots (**a**) or *M. polymorpha* thalli (**b**) compared to axenic culture. Significantly differentially expressed genes (DEGs) are in red. **c,** Hierarchical clustering of differentially expressed secreted protein genes of Fol during colonization of tomato *S. lycopersicum* (Sl) or *M. polymorpha* (adjusted p ≤ 0.05; log2 fold change [logFC] ≥ 2) at 3 dpi. Colour scale indicates expression level. **d-e,** Volcano plots showing pairwise differential expression analysis of Fol secreted protein genes encoded on core (**d**) and lineage-specific (**e**) genomic regions during infection of tomato (left) versus *M. polymorpha* (right) at 3dpi. Green = 2 < Log2FC < 8; Red = 8 < Log2FC < 15 and Blue = Log2FC > 15. Functionally characterized SIX and ERC effectors are highlighted.

We identified a set of 500 and 421 Fol genes encoding predicted secreted proteins that were significantly upregulated (log2FC > 2, padj < 0.05) during colonization of Mp or tomato respectively, accounting for 27.7 % and 23.3 % of the total secretome of Fol expressed at the infection time point tested (Figure. 2c). A chromomap of the secreted protein genes induced during infection of Mp or tomato revealed that 89.4 % and 85.7 % respectively, were located on core genomic (Supplemental Figure. 3a,b). This indicates that the establishment of early compatibility^15^ of this vascular wilt pathogen with phylogenetically distant plant hosts is primarily mediated by core-genomic regions.

**Figure. 3.**
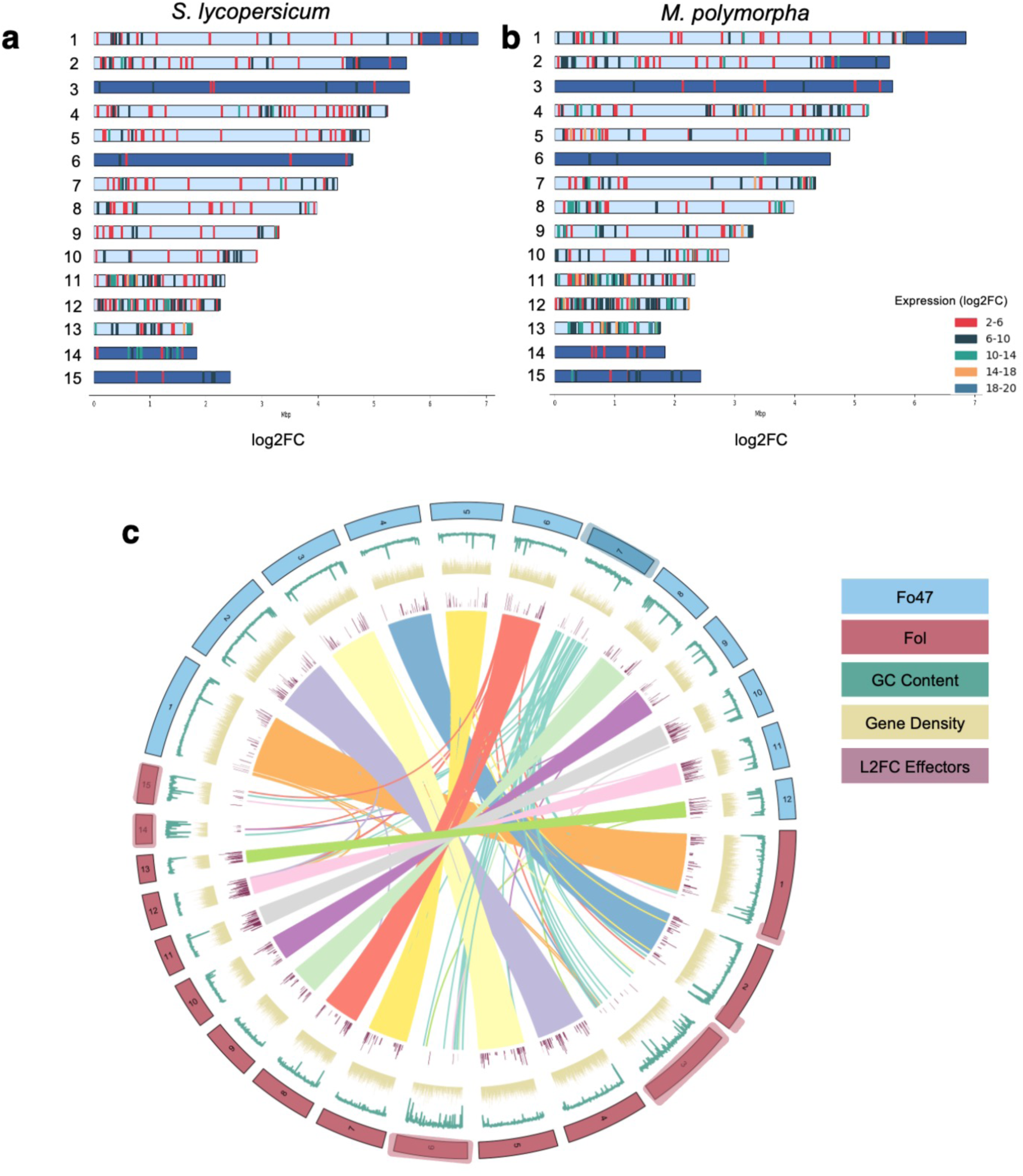
Genes encoded on fast core chromosomes drive plant associations of *F. oxysporum*. **a-b,** Chromosomal distribution of Fol genes encoding predicted secreted proteins upregulated during infection of tomato (**a**) or *M. polymorpha* (**b**). Each bar represents a gene on the chromosome with colour indicative of expression (log2FC). Core regions of the genome are indicated in light blue and lineage-specific regions are shown in navy blue **c,** Circos plot showing synteny between genome assemblies of *F. oxysporum* isolates Fol and Fo47. Blue/Red, Karyotypes of assembled chromosomes; Green, GC content (%); Yellow, Gene density, both calculated in 10 kb windows; Mauve, Log2FC of *in planta* upregulated genes encoding predicted secreted proteins, for the sake of visibility, gene -start and end- were multiplied by 10 to increase the effective gene length; Syntenic relationships shown by linking syntenic gene blocks (single copy orthogroups) in each genome pair. Core chromosomes can be identified through synteny between Fol and Fo47, whereas accessory chromosomes show no or reduced synteny. Lineage-specific regions are highlighted.

Importantly, the *in planta*-induced genes encoding predicted secreted proteins included the previously characterized core effectors ERC1, ERC3 and ERC4 [15] as well as the lineage-specific SIX effectors SIX1, SIX3, SIX5 and SIX6 [23–25] (Figure. 2d,e). However, most of the secreted protein genes upregulated during Mp colonization were located on core genomic regions, while the secreted protein encoding genes upregulated during colonization of tomato roots were located both on core and LS regions (Figure. 2d,e). Furthermore, *ERC* effector genes were induced during colonization of both Mp and tomato at comparable levels, while upregulation of *SIX1*, *SIX3*, *SIX5*, *SIX6* was only detected on tomato, with relative expression levels 5 to 6 fold higher compared to Mp. Analysis of the expression profiles of four effector candidates, two encoded on a fast core and the other two on a LS chromosomes, showed a similar expression pattern of the fast core effector on both plant hosts, with a higher induction in comparison to the LS effector candidate (Supplemental Figure. 4a-d). Collectively, our analysis reveals a tight transcriptional control of secreted proteins encoded on core and LS on the different plant hosts. These results corroborate the previous finding that the basal ability of Fo to infect a non-vascular plant host is independent of the host-specific effectors since isogenic mutants lacking LS effectors show no detectable differences in disease severity on Mp, but are reduced in virulence on a vascular plant host [21,24,25–27].

**Figure. 4.**
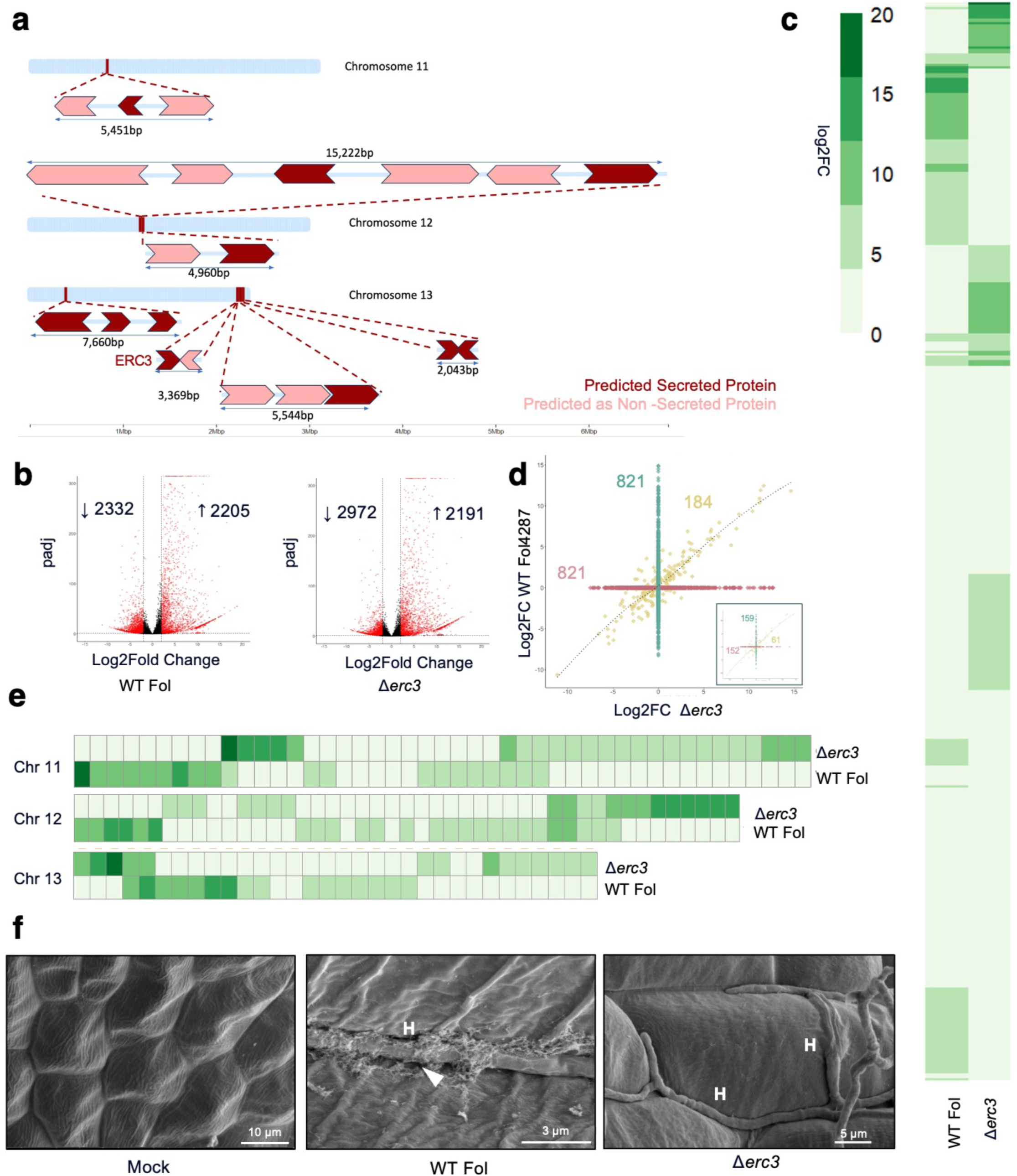
Gene clusters located on fast core chromosomes are upregulated during colonization of phylogenetically distant hosts. **a,** Schematic representation of the genomic regions on Fol chromosomes 11, 12 and 13 harbouring gene clusters that are transcriptionally activated during colonisation of tomato and *M. polymorpha*. Genes are depicted as arrows. Dark and light colours indicate secreted and non-secreted proteins, respectively. The previously characterized *ERC3* effector gene is indicated. **b-c,** Volcano plot showing pairwise differential gene expression analysis in the Fol wild type (**b**) or Δ*erc3* mutant (**c**) during infection of *M. polymorpha* at 3dpi, compared to axenic culture. Significantly DEGs are in red. **c,** Hierarchical clustering of secreted protein genes in Fol wild type or Δ*erc3* mutant during infection of *M. polymorpha* Tak-1 (adjusted p ≤ 0.05; log2 fold change [logFC] ≥ 2) at 3 dpi. Colour scale indicates expression level. **d,** Scatter plot representing the expression of secreted protein genes from Fol wildtype or Δ*erc3* mutant. Green and red are genes expressed exclusively in wildtype or Δ*erc3* mutant, respectively, while yellow represent genes upregulated in both. Inset shows the same information for genes located on fast core chromosomes. **e,** Comparative heatmap showing expression of secreted protein genes located on fast core chromosomes in the Fol wildtype or Δ*erc3* mutant during infection of *M. polymorpha*. **f,** SEM micrographs of *M. polymorpha* thalli infected with the indicated strains at 3 dpi. Note hyphae (H) entering the thallus surface, deposition of fibrous material in the wildtype likely representing remnants of degraded host cell wall (arrow) and its absence in the Δ*erc3* mutant. Scale bars = 10 µm in left image; 3 µm in middle image; 5 µm in right image.

### Fast core chromosomes encode genes activated during associations with evolutionarily distant plant hosts

Fo has a compartmentalized genome that contributes to the higher adaptability of this fungus. A ‘division of labor’ between genomic regions has been suggested where LS regions determine host-specific pathogenicity, while core regions contribute to generic infection-related processes [28]. The three smallest core chromosomes of Fol – chromosomes 11, 12 and 13 have been called ‘fast-core’ chromosomes since they are more divergent in terms of sequence similarity and synteny as compared to the larger core chromosomes [28]. Despite having a similar gene density as the core chromosomes, the fast core chromosomes encode genes that tend to be upregulated during infection and are enriched in the H3K27me3 methylation marks which occupied ∼93% of the fast core chromosomes [12,28].

Here we found that the fast core chromosomes 11, 12 and 13 were particularly enriched in the secreted protein genes upregulated during colonization of the vascular and the non-vascular plant host (Figure. 3a,b and Supplemental Figure. 3a). Of the total DEGs of Fol upregulated at the infected timepoint tested, 465 (21.7% in Tomato) and 647 (23.4% in Mp) were located on the fast core chromosomes, with 25.5% and 28.6% respectively, encoding small secreted proteins representing potential effectors (Supplementary Table. 1). Furthermore, most of the genes upregulated >10-fold during infection of either host were located on a fast core chromosomes (Figure. 3a,b; Supplementary Table. 1). Functional annotation with InterPro of potential domains encoded by these upregulated secreted genes showed an enriched function in Glycoside hydrolase, cellulose binding and peptidase biological processes (Supplementary Table. 2). We tested whether *in planta* induction of genes encoded on fast core chromosomes is conserved in a Fo isolate enhibiting an endophytic lifestyle by performing RNA-seq of Mp Tak-1 thalli infected with Fo47 at 3 dpi. This identified 465 putative secreted protein genes of Fo47 that were significantly upregulated on Mp (log2FC > 2, padj < 0.05) (Supplemental Figure. 3b,c; Supplementary Table 1). A chromomap of these differentially expressed secreted genes showed that around 37% were located on fast core chromosomes (Supplemental Figure. 3b; Supplementary Table 1). Again, most genes upregulated >10-fold in endophytic isolate Fo47 were located on a fast core chromosome (Supplemental Figure. 3b).

CAZyme profiling of Fol and Fo47 genes induced during Mp infection revealed differences in the percentage as well as in the absolute number of CAZyme families between Fol versus Fo47 (Supplemental Figure. 5a,b). Enrichment analysis using the Chi-Square mosaic plot revealed a significantly higher probability of an upregulated gene to be located on a fast core chromosome for Fol on Tomato (!^2^ = 8.998, p = 0.002); or for Fol (!^2^ = 398.35, p = 2.22e-16) or for Fo47 (!^2^ = 287.45, p = 2.22e-16) on Mp (Supplemental Figure. 6a–c). Collectively, these results suggest that secreted protein genes encoded on fast core chromosomes drive Fo associations with evolutionarily distant land plants as well as across different lifestyles reaching from endophytic to pathogenic.

**Figure. 5.**
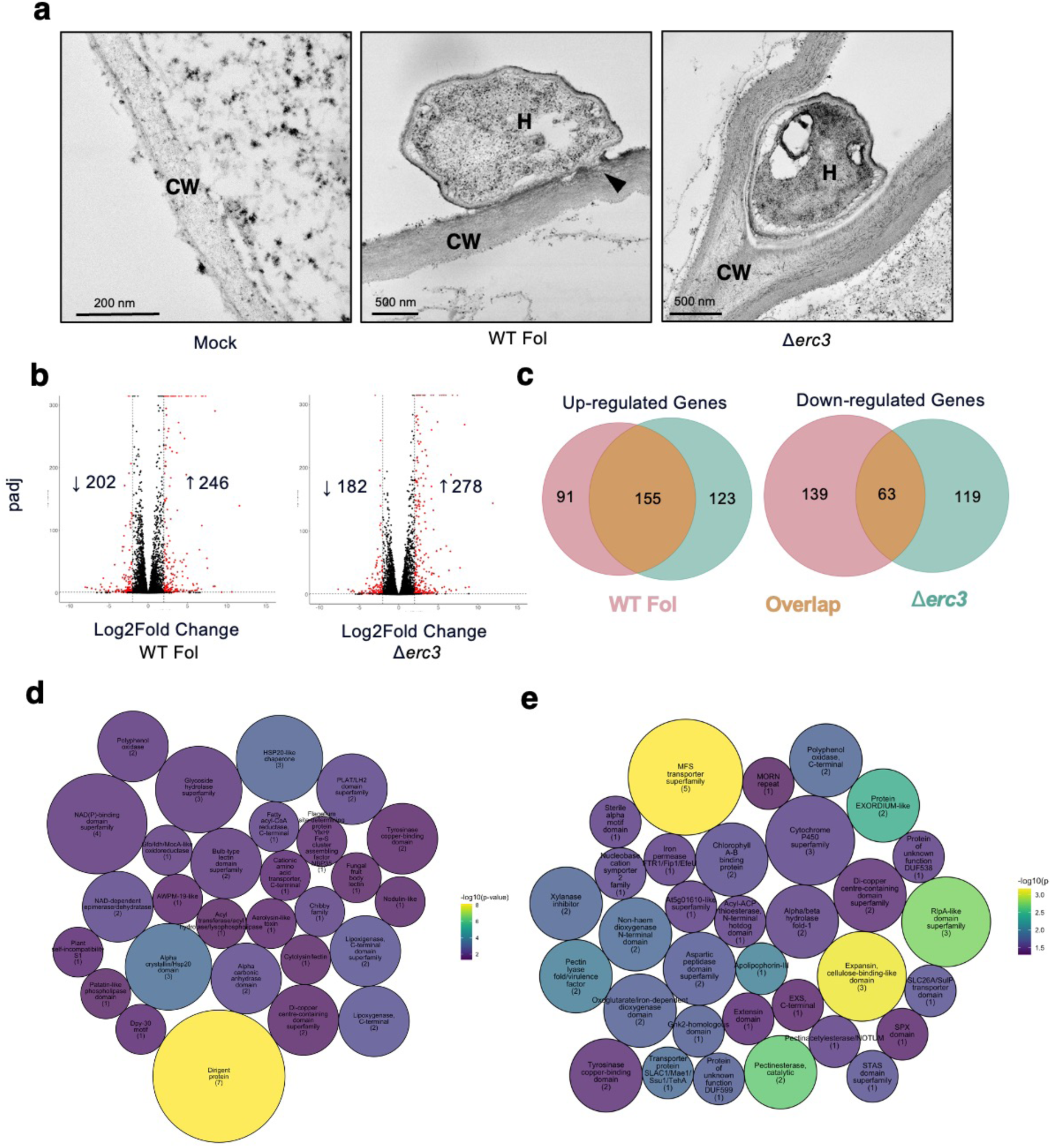
Deletion of *ERC3* leads to enhanced cell-wall mediated defence responses in *M. polymorpha*. **a,** TEM micrographs of *M. polymorpha* thalli infected with Fol wildtype or Δ*erc3* mutant at 3 dpi. Note intact cell wall (CW) in the uninfected control, in comparison to the infected thallus showing hyphae (H) of the wildtype attempting to breach the plant cell wall and entering the thallus cell. Hyphal penetration leads to death of the cell which is devoid of cytoplasm (arrowhead). In contrast, during colonization of the Δ*erc3* mutant, fortification of the host cell wall is observed with hyphae growing intercellularly. Scale bars = 200 nm in left image; 500 nm in middle and right image. **b,** Volcano plot showing pairwise differential expression analysis of *M. polymorpha* genes upon infection with Fol wildtype or Δ*erc3* mutant at 3 dpi. **c,** Venn diagram showing overlap of the transcriptomes from (**b**). **d-e,** Interpro Term Enrichment analysis using FUNC-E package (ecut <0.05) on genes downregulated in *Marchantia* show enrichment of dirigent proteins indicating an active immune suppression by Fol wildtype (**d**) but not by Δ*erc3* mutant (**e**). The sizes correspond to the number of genes in each IPR term, and the statistical significance is represented as a -10 log of Fisher’s p-value. A custom R script was written using the pack circles and the ggplot package was used to visualize the enrichment.

**Figure. 6.**
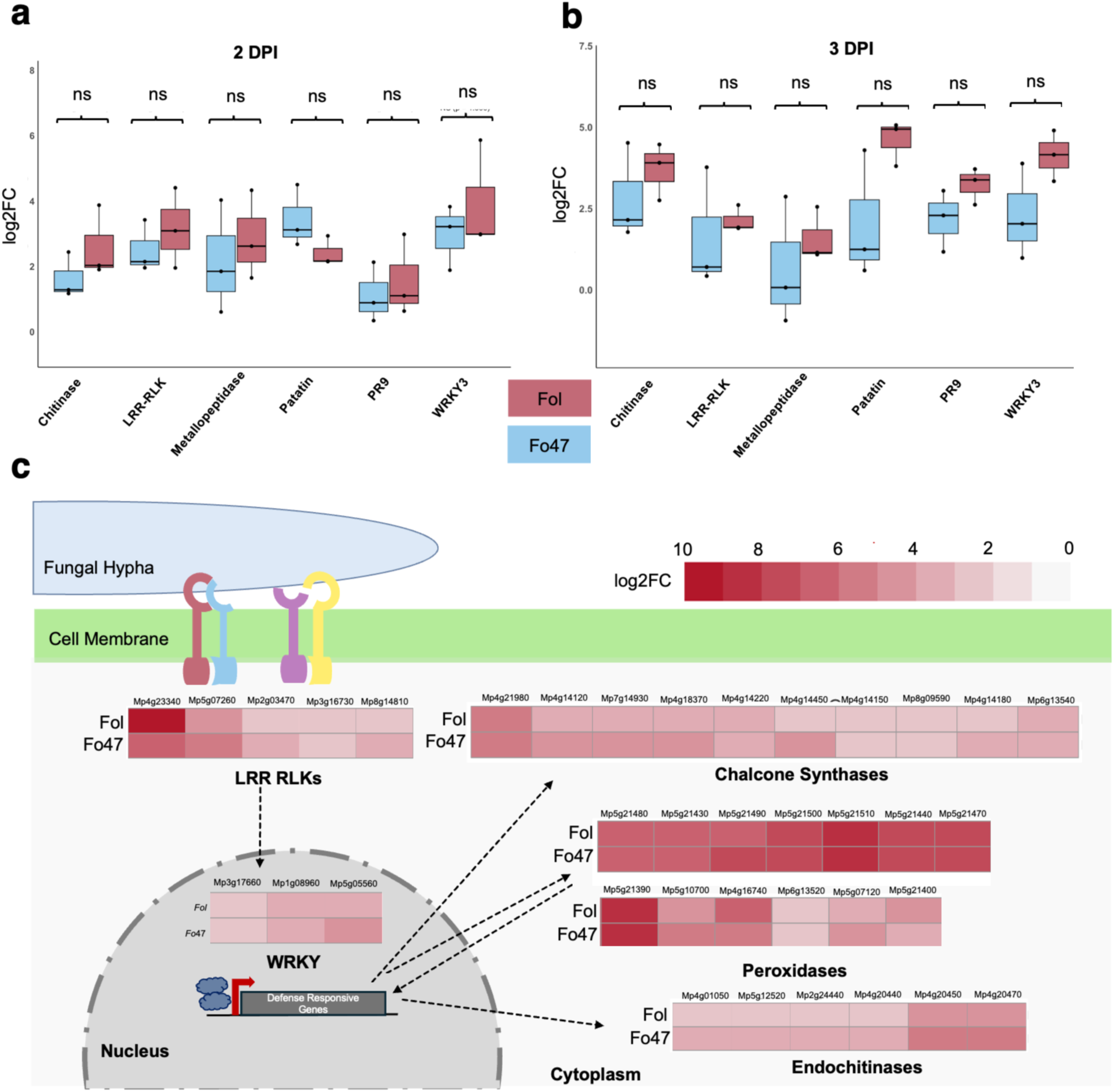
*M. polymorpha* mounts a general transcriptional response against different *F. oxysporum* isolates. **a-b,** Relative transcript levels of defense genes in *M. polymorpha* upon infection with the *F. oxysporum* isolates Fol and Fo47. Transcript levels of the selected genes were measured by RT-qPCR of cDNA obtained from thalli either uninoculated or inoculated with either fungal isolate at 2 (**a**) or 3 dpi (**b**). Values shown are the average of three biological replicates with technical replicates each, calculated as log2(2^-ΔΔCt^) using the *MpEF1*α gene as internal control. Lines and boxes represent the arithmetic mean and the interquartile range, respectively. Significance was calculated using Mann-Whitney U Test with an asymptotic normal approximation of the U statistic. NS = non-significant. **c,** The conserved response of *M. polymorpha* to Fol and Fo47 represented as log2FC inferred from bulk transcriptomics and represented as a schematic overlay of a cell.

### Fast core chromosomes show conserved effector profiles across Fo isolates

Comparative genomics was used to investigate synteny in the genomes of Fol and Fo47, which display contrasting lifestyles on Tomato. OrthoFinder [29] identified single-copy orthologous protein groups between the two tested isolates and visualised the synteny between genomes (Figure. 3c). Furthermore, we calculated the GC content and gene density across chromosomes in a sliding window of 10 kb to validate the LS regions in the two genomes. We observed a clear demarcation of LS regions with high GC content and low gene densities in chromosomes 3, 6, 14 and 15 of Fol, along with some terminal regions of chromosomes 1 and 2 as reported previously [12]. We further detected chromosome 7 of Fo47 as being LS^30^. We confirmed the conservation of fast core chromosomes 11, 12, and 13 of Fol with their homologous chromosomes 10, 11, and 12 of Fo47 (Figure. 3c). Representation of L2FC secreted genes across these chromosomes revealed a conserved effector catalogue upregulated *in planta* specifically on the fast core chromosomes, across both Fo isolates (Figure. 3c). The three-speed evolutionary hypothesis also proposes the fast core chromosomes to evolve at an intermediate rate between core and LS regions [28]. Considering this, our results suggest that the fast core chromosomes, exhibiting a similar expression profile of genes upregulated during Mp infection, potentially play a role in establishment of basic compatibility of Fo isolates across evolutionarily distant plant lineages.

### Gene clusters located on fast core chromosomes are transcriptionally activated during infection of non-vascular and vascular plant hosts

Filamentous plant pathogens have an uneven distribution of genes and repetitive elements across and between chromomosomes [12,31–34]. In plant pathogenic fungi, genes encoding metabolic enzymes often occur in co-expressed clusters [35,36]. Furthermore, gene clusters encoding secreted protein effectors have been described in the corn smut fungus *Ustilago maydis*, where genes encoding secreted proteins upregulated during tumor formation are arranged in 12 clusters comprising of 3 to 26 genes [37]. Deletion of several gene clusters in *U. maydis* evidenced their role in virulence on maize [37–39]. Given the previously suggested role of Fo genes on fast core chromosomes during infection and in metabolism, transport or defense [28], we determined the genomic positions of the *in planta* upregulated genes located on fast core chromosomes and aimed to identify pairs or clusters of genes which are induced during colonization.

Gene clusters were identified following the criteria previously defined for *pit* cluster proteins in *U. maydis* [38], where neighbouring genes are less than 2 kb apart and at least one of the genes is predicted to be secreted. This identified 12 and 11 putative gene clusters on fast core chromosomes that were upregulated during infection of Fol on Mp or tomato, respectively (Supplementary Table. 3). Six of these clusters were shared across both the hosts (Figure. 4a), and five of them also showed upregulation in the endophytic strain Fo47 during Mp colonization (Supplemental Figure. 7a, Supplementary Table. 4). The five shared clusters upregulated in both Fo isolates and on both plant hosts comprises between 2 and 6 genes.

While some genes encode hypothetical proteins, others code for a transmembrane protein or show homology with pectate lyase and glycoside hydrolase domain proteins (Supplementary Table. 5). Interestingly, two of these shared clusters on chromosome 13 encode uncharacterized hypothetical secreted proteins which are upregulated 3 to 14-fold during colonization of Mp or tomato. Almost all Fo genes located in these shared clusters showed 2-18 fold upregulation during infection of both plant hosts (Supplementary Table. 3). Importantly, the previously identified core effector *Erc3* (<u>E</u>arly <u>R</u>oot <u>C</u>olonization 3; FOXG_16902) together with FOXG_16903 in a head-tail orientation, constituted one of the shared gene cluster pairs located on chromosome 13 (Figure. 4a, Supplemental Figure. 7a). ERC3 was identified during intercellular colonization of tomato roots and contributes to Fol virulence [15]. FOXG_16903 encodes a hypothetical protein with twelve predicted transmembrane domains [40] (Deep TMHMM; Supplemental Figure. 7b). This arrangement is reminiscent of the *pit* gene cluster of *U. maydis* where Um*Pit1* encodes a transmembrane protein and *UmPit2* encodes a secreted protein that contributes to corn smut [38]. Together, these findings suggests that gene clusters on fast core chromosomes are actively transcribed during infection of phylogenetically distant plant hosts as well as across Fo isolates exhibiting contrasting lifestyles on angiosperms.

### Deletion of *erc3* leads to misregulation of predicted secreted protein genes and impairs host colonization

In light of the previous observation that FoERC3 contributes to virulence on Mp [15], we asked whether deletion of *erc3* affects the dynamics of infection. Transcriptomic profiling of Fo was performed by bulk RNA-seq of Mp Tak-1 thalli infected either with wild-type Fol or the Δ*erc3* mutant at 3 dpi, and compared to axenic fungal culture or mock infected thalli. This revealed 4,537 and 5,163 Fo genes that differentially expressed between wildtype Fol or the Δ*erc3* mutant during colonization of Mp (Figure. 4b, Supplemental Figure. 8a,b Supplementary Table. 6).

Profiling of genes encoding predicted secreted proteins expressed differentially between Fol and Δ*erc3* (Fig. 4c) identified around 1005 secreted protein encoding genes (log2FC > 2, padj < 0.05) with a distinct expression pattern, including 821 genes unique for either isolate compared to only 184 genes shared among both (Figure. 4d). These results indicates a strong transcriptional rewiring of the secretome upon *erc3* deletion. Kendall’s correlation test for the 184 genes expressed in both isolates revealed a moderately positive correlation (“= 0.6914944, p-value < 2.2e-16) represented as overall trend with a LOESS curve (Locally Estimated Scatterplot Smoothing). This pattern remained relatively constant when we zoomed in on the fast core chromosomes, indicating that alteration of the secretome is general and not limited to the fast core chromosomes (Figure. 4d, inset and Figure. 4e).

A misregulation of the secreted protein genes was detected on fast core chromosomes 11 to 13. Around 83.6% of the secreted protein genes encoded on fast core chromosomes showed an exclusive differential expression either in Fol or the Δ*erc3* mutant. In addition, 42% of the 61 secreted protein genes encoded on the fast core chromosomes that are expressed in both Fol wildtype and Δ*erc3* showed higher expression in the Δ*erc3* mutant, indicating a possible compensatory effect in the Δ*erc3* mutant. Previous studies suggested that ERC3 is a putative %-N-arabinofuranosidase B, which cleaves oligosaccharides to release arabinose residues [15].

We therefore investigated the ultrastructure of the Δ*erc3* mutant during Mp colonization. SEM showed Fol wildtype hyphae to deposit a reticulate material (likely originating from the host cell) at the site of entry during intercellular colonization of the host, while no such components were observed in Δ*erc3* mutant (Figure. 4f). On the contrary, the Δ*erc3* mutant remained largely restricted to the intercellular space and failed to enter the host cells (Figure. 4f). Overall, these results suggest that disruption of a gene located in a cluster on a fast core chromosome leads to misregulation of the transcriptional program and compromises intracellular colonization of host.

### Cell wall-mediated defense responses in Mp are enhanced during interaction with the *Δerc3* mutant

ERC3 is a predicted %-N-arabinofuranosidase B, which can cleave oligosaccharides to release arabinose during host-mediated cell wall remodeling. Previous work in the rice blast fungus *Magnaporthe oryzae* showed that *Mo*Abf, an %-L-arabinofuranosidase B, causes cell wall degradation in rice leaves triggering a host immune response [41]. Although FolERC3 shares only 23.69% amino acid sequence identity with MoAbf, both carry the characteristic AbfB domain predicted by Interproscan. We used Transmission Electron Microscopy (TEM) to study plant cell wall penetration during Mp colonization at the ultrastructural level. The Fol wildtype strain was able to breach the host cell wall (Figure. 5a, arrowhead in the middle panel) in multiple locations, whereas the Δ*erc3* mutant did not penetrate the host cell. Instead, the hyphae remained trapped within the intercellular space, with signs of host cell wall fortification (Figure. 5a). Furthermore, penetration of the cell wall by wildtype hyphae led to death of the host cell, detected as a loss of electron density compared to the uninoculated host. By contrast, minimal cell death signatures were detected in the Δ*erc3* infected tissue (Figure. 5a) likely due to the lack of intracellular penetration.

Next, we performed a transcriptomic analysis by bulk RNA-seq of Mp thalli infected with Fol wildtype and Δ*erc3* at 3 dpi. Pairwise differential gene expression analysis detected with around 450 DEGs across the two fungal isolates testes (Figure. 5b and Supplementary Table. 7). A 42% overlap was observed for the DEGs upregulated in Mp infected by the Fol wildtype vs. Δ*erc3*, while the overlap in the downregulated DEGs of Mp was only 19.6% (Figure. 5c). This suggests that although Mp perceives the two isolates in a similar way, Δ*erc3* fails to suppress part of the host defense response. To further elucidate these differentially activated plant pathways, we performed InterPro Term Enrichment analysis to identify functional or structural protein motifs enriched in our dataset of Mp genes upregulated in infection by Δ*erc3* but not be the Fol wildtype. FUNC-e package for Python was used to enrich at the cutoff of e <0.001. This revealed an enrichment of plant cupredoxins, riboflavin-synthases, FAD binding, multicopper oxidases, etc, as common responses activated in Mp by both isolates. By contrast, chitin-binding/endochitinase-like, glucosyl hydrolase superfamily genes and dirigent proteins were enriched exclusively in Mp DEGs either upregulatred or downregulated during infection with the Fol wildtype only, whereas the glycosidase superfamily was enriched among Mp genes upregulated during infection by the Δ*erc3* mutant (Supplemental Figure. 8c,d; Figure. 5d,e). These findings indicate that the wildtype Fol is able to penetrate the host cells whereas the Δ*erc3* mutant is not. Instead, it triggers a cell wall-mediated plant defense response, thereby

preventing the fungal hyphae from intracellular host colonization. Our results also suggests that the cell wall mediated defense responses upon detection of filamentous pathogens are conserved across evolutionarily distant land plants.

### Mp mounts a generic transcriptional response against Fo isolates and other pathogens

Isolates of Fo can trigger a pathogen associated molecular pattern (PAMP) response in Mp by activating PAMP responsive genes [21]. We hypothesized that if conserved secreted proteins encoded on fast core chromosomes drive colonisation of Mp by different Fo isolates, the host immune response to these isolates should be similar. To test this we measured expression of a panel of Mp homologs of plant defense genes representing different steps of pattern-triggered and effector-triggered immunity (PTI & ETI) in angiosperms. These include chitinase, leucine-rich repeat receptor kinase (LRR-RLK), metallopeptidase, patatin PR9 and the transcription factor *WRKY3*. Expression profiling of these defense marker genes in Mp thalli infected either with Fol or Fo47 at 2 and 3 dpi failed to detect significant differences between the two Fo isolates (Figure. 6a,b). This indicated the presence of a common defense response in Mp against different Fo isolates.

We next compared the transcriptional profiles of Mp thalli inoculated with Fol or Fo47 in comparison to the water control. Differential expression analysis (log2FC >2 and padj <0.05) revealed a similar transcriptional profile of Mp in response to Fol and Fo47 infection, with around 301 (Fol) and 573 (Fo47) DEGs, 57% of which were induced by both fungal isolates (Supplementary Table 8). Using Interproscan, we predicted the domains in Mp proteome (Supplemental Table 9). In the set of genes upregulated (log2FC >2) in our dataset of Fol and Fo47 infected thalli, we identified genes encoding for proteins involved in defense responses and plotted their expression on a schematic overlay of a cell. Genes encoding for LRR-RLKs (e.g. *Mp4g23340*) were upregulated around 7 and 10 fold upon Fo47 and Fol infection, respectively. Furthermore, WRKY transcription factor genes such as *WRKY3*, *WRKY6*, and *WRKY7* were also induced in response to both fungal isolates (Figure. 6c). Finally, chalcone synthase functioning in the flavonoid and anthocyanin synthesis pathway as well as a dozen peroxidases and endochitinases were also significantly enriched in response to both Fo isolates. All these Mp genes exhibited similar levels of expression in response to the two fungal isolates, pointing towards a common transcriptional response of Mp to Fo.

We next tested whether the general response to different Fo isolates is also activated in response to other Mp pathogens such as the oomycete *Phytophthora palmivora* [19], the bacterial pathogen *Pseudomonas syringae* DC3000 [20] or the fungal pathogen *Colletotrichum nymphaeae* [42]. Among the core defense gene, most of the commonly upregulated gene families such as chalcone synthase, endochitinase, glycosyl hydrolase, peroxidase, LRR-RLK and WRKY were also induced in response to these other pathogens with a similar expression profile, suggesting a common defense activation mode against microbial invaders (Supplementary Table. 10). Overall, these results suggest that Mp mounts a general immune response to different microbes including those exhibiting endophytic and pathogenic lifestyles on angiosperms.

## DISCUSSION

The availability of complete genome sequences and the establishment of different biotic interaction models in the liverwort Mp have made this the main model of study for evoMPMI and for understanding conserved host mechanisms that are exploited by microbes to establish plant associations [1,18,19,21,43]. During the last few years, studies have mainly focused on understanding the core immune mechanisms across land plant lineages [44,45]. However, the mechanisms of microbial pathogenicity to colonize distinct land plants remain underexplored. Here we investigated the crosstalk during infection of Mp by different isolates of the vascular wilt fungus Fo, which display contrasting lifestyles – pathogenic vs. endophytic - on flowering plants. Our goal was to monitor the transcriptional response of Fo across the distinct genomic regions - core vs. LS – during infection of two phylogenetically distant land plants.

Previous findings indicated that the core pathogenicity mechanisms are sufficient for the infection of the non-vascular host Mp, while and LS regions confer host-specific vascular adaptation on angiosperms [21]. Based on these observations, we hypothesized that Fo association with distinct plant lineages is based on regulation of genes encoded on core genomic regions, which are conserved across the Fo species complex. Transcriptomic analysis of the expression profiles of core and LS effectors at 3 dpi, revealed upregulation of secreted protein genes encoded on the core genomic regions, including the conserved ERC effectors, both on Mp and on the vascular host tomato. In contrast, the secreted protein effectors encoded on the LS regions, including the SIX effectors were specifically upregulated during infection of Tomato as compared to Mp. This indicated that the latter are largely dispensable for infection of a non-vascular host and points towards a differential transcriptional control of core and LS virulence effectors during establishment on vascular and non-vascular plant lineages. The basal ability of Fo to establish compatibility on land plants appears to be determined by the broadly conserved core regions. Interestingly, the partial upregulation of some of the *six* genes during infection of Mp suggests a possible role independent of the xylem-specific functions reported previously [46].

We detected an enrichment on fast core chromosomes of upregulated secreted protein genes in different Fo isolates with contrasting lifestyles, during infection of flowering and non-flowering plants. These results are consistent with previous reports suggesting that fast core chromosomes are enriched for genes upregulated during infection [28]. This work also suggested that genes on fast core chromosomes have lower expression levels compared to those from the core genome, both *in vitro* and *in planta* [28]. However, under the conditions tested here the secreted protein genes encoded on fast core chromosomes displayed an induction of up to 18-20 fold Log2FC. These data suggest that genes encoded on fast core chromosomes drive plant associations across evolutionarily distant land plant lineages in different Fo isolates. This is in line with the previous finding that transcriptional waves of effector genes encoded on distinct genomic regions modulate lifestyle transitions of Fol from endophytic to pathogenic [15]. Interestingly, the fast core chromosomes of different Fo isolates were highly syntenic for transcriptionally activated effector catalogues, which showed similar expression profiles during infection of Mp. These results point towards a genetic program for plant association encoded by fast-core chromosomes.

Our analysis identified seven clusters containing predicted secreted protein genes that were upregulated on phylogenetically distant hosts and in Fo isolated with distinct lifestyles. Interestingly similar clusters of effector genes have been reported to contribute to virulence in other fungal phytopathogens [37]. In *U. maydis* a neighboring gene pair encoding a transmembrane and a secreted protein was co-regulated during infection and required for virulence [38]. We found that clusters of genes conserved in two Fo isolates were upregulated 18-fold during infection of a non-vascular and a vascular plant. One upregulated gene pair showing a head to tail orientation contains the previously characterized core effector ERC3. Disruption of Δ*erc3* led to misregulation of multiple secreted protein genes encoded on fast core chromosomes. Interestingly, some of these showed increased transcript levels in the Δ*erc3* mutant. Collectively, our findings suggest that transcriptional plasticity of gene clusters on fast core chromosomes of Fo encoding secreted proteins drives colonization of land plants. Further characterization of these individual gene clusters will be crucial to uncover their precise role during establishment of host compatibility.

One application of the Mp model in EvoMPMI is to understand conserved or lineage-specific plant immune mechanisms and how these plant immune networks evolved. The recent development of the Mp pangenome revealed ancient mechanisms of plant adaptation, including peroxidases or nucleotide-binding leucine-rich repeats (NLRs) that are conserved in angiosperms [47]. We observed a general transcriptional response of Mp to different isolates of Fo, including a core set of plant defense genes including LRR-RLKs, WRKY transcription factors, chalcone synthase, peroxidases and endochitinases, which showed a similar induction upon inoculation with Fol or Fo47. Similar responses were detected upon biotic challenge with *Phytopthora palmivora* [19] and *Pseudomonas syringae* DC3000 [20]. This suggests the presence of a general defense program in Mp, with a core set of gene families in response to different microbes.

Taken together, our findings suggest a role of modular genome organization and its transcriptional regulation during establishment of broad-spectrum plant association in Fo. Mechanisms encoded on fast core chromosomes appear to drive fungal establishment during the asymptomatic phase both on vascular and non-vascular hosts. Future studies should provide new insights into the mechanisms of fungal adaptation from basal plant association to niche specialization leading to some of the most destructive systemic diseases on angiosperms.

## MATERIALS AND METHODS

### Fungal strains and growth conditions

This study utilized *F. oxysporum* isolates, including the tomato-pathogenic strain Fo f. sp. *lycopersici* and its derived Δ*erc3* (FOXG_16902) knockout mutant [15], the endophytic isolate Fo47. Fungal cultures, preparation of the spore inoculum and their glycerol stocks were performed as described [48].

### Strains and growth conditions

Fo isolates were grown in Potato Dextrose Broth (PDB) for 72 hours at 28°C and 140 rpm. Microconidia were harvested by filtering fungal cultures through the cheese cloth. The spores were pelleted by centrifugation and resuspended in sterile water [49]. Spore suspension for infection was created by diluting the spore suspension to a final concentration of 10^5^ spores/mL.

### Growth conditions and infection of Mp

Gemmae from the Mp Tak-1 were plated on Gamborg B5 media without sucrose, solidified with 1.5% agar, and maintained at 22°C under a 20:4-hour day-night photoperiod. Growth and maintainance of Mp was done as described previously [50]. For infection, 21-day-old thalli (≥ nine thalli per treatment) were dipped in microconidial suspension for 20 mins. Dipping in water was performed as a mock/control treatment. After air drying for 15 minutes, the infected thalli were incubated at 28°C with a 20:4-hour day-night cycle and samples were harvested at the defined time points.

### Transmission and scanning electron microscopy

Small pieces (1 mm^2^) of leaves were fixed in 2.5% glutardialdehyde dissolved in 0.06 M phosphate buffer (pH 7.2) for 90 min. Samples were then washed in buffer (4 times 15 min) and post fixed with 1% osmium tetroxide dissolved in 0.1 M phosphate buffer (pH 7.2) for 90 min. After a rinse in buffer (4 times 10 min) samples were dehydrated in increasing concentrations of acetone (50, 70, 90, and 100%) for 2 times 10 min each. Samples were then separated in two groups: 1.) For TEM-investigations one group was infiltrated with increasing concentrations (30%, 50%, 70%, and 100%) of epoxy resin (EMbed 812; Electron Microscopy Sciences, Hatfield, PA, USA) mixed with acetone for at least 1 h each step. Polymerization was performed with pure EMBed 812 resin at 60 °C for 48 h. Ultra-thin sections (80 nm) sectioned with an EM UC7 ultramicrotome (Leica Microsystems, Buffalo Grove, IL) were post-stained with lead citrate (1% dissolved in 0.6 M NaOH) for 5 min and uranyl-acetate (2% dissolved in aqua bidest) for 15 min. Sections were imaged with a Spectra 300C TEM (Thermo Fisher Scientific, Hillsboro, OR). 2.) For SEM-investigations the other group of samples was critical point dried (Leica EM CPD 300; Leica Microsystems) using a customized program for plant leaves with a duration of 80 min (settings for CO2 inlet: speed = medium & delay = 120 s; settings for exchange: speed = 5 & cycles = 18; settings for gas release: heat = medium & speed = medium). Dried samples were mounted on aluminum stubs with carbon tape, sputter-coated with 10 nm iridium (Leica EM ACE 600; Leica Microsystems) and imaged with a Versa 3D scanning electron microscope (Thermo Fisher Scientific, Hillsboro, OR).

### Quantification of fungal biomass

qPCR was performed on genomic DNA extracted from 3 dpi infected thalli to quantify fungal biomass. DNA extraction was carried out using the 2X CTAB method. Briefly, the tissue was snap-frozen liquid nitrogen and crushed in an extraction buffer (1.4% SDS, 100 mM Tris-HCl, pH 8, 0.5M NaCl and 0.0072% v/v β-mercaptoethanol). An equal volume of 2X CTAB buffer (2% CTAB, 100 mM Tris-HCl, pH 8, 1.4 M NaCl) was added, and the tubes were incubated at 65°C for 10 minutes. Phase separation was carried out using 24:1 Chloroform: isoamyl alcohol, and DNA was precipitated in isopropyl alcohol. After washing the DNA pellet with 70% ethanol, the DNA was resuspended in sterile water. Fungal biomass was quantified using qPCR with 50 ng DNA in each well using the iTaq Universal SYBR Green Supermix and 0.2 µM of forward and reverse primers (complete list of oligos in Supplementary Table 11). Mp*actin* (Mp6g11010) and Fo*actin* (FOXG_01569) were used as plant and fungal DNA readouts, respectively. The PCR was carried out with an initial denaturation at 95°C for 10 minutes, followed by 40 cycles of 95°C for 10 seconds, 60°C 10 seconds, and 72°C for 20 seconds. Fungal biomass was represented as the a log2 ratio of Ct*Mpactin* and Ct*FoActin*.

### RNA extraction and Gene Expression Analyses using RT-qPCR

RNA was extracted from snap-frozen thalli at defined time points using RNA extraction was performed using Qiagen RNeasy Kit for RNA Purification, using 100 mg tissue and 500 µL of lysis buffer (washing and elution were done as per the manufacturer’s guidelines). RNA was eluted in 40 µL nuclease water, followed by DNase treatment. cDNA synthesis was carried out using PrimeScript™ 1st strand cDNA Synthesis Kit. qPCR was performed with the iTaq Universal SYBR Green Supermix and 0.2 µM of forward and reverse primers (list of oligos in Supplementary Table. 11). The PCR was performed with an initial denaturation at 95°C for 10 minutes, followed by 40 cycles of 95°C for 10 seconds, 60°C 10 seconds, and 72°C for 20 seconds. *FoActin* and Mp*EF1α* (Mp3g23400) were used as an internal control for profiling of fungal and *Marchantia* genes, respectively. Gene expression was calculated using the ΔΔCt method [51].

### Library preparation & sequencing

The extracted RNA from 3dpi infected thalli was eluted in 40 µL nuclease-free water and quantified using Qubit and quality assessment was performed on Agilent TapeStation. From the total RNA, mRNA was enriched using NEBNext Poly(A) mRNA magnetic isolation Module. The library was prepared using NEBNext Ultra™ II Directional RNA Library Prep with sample purification beads. Sequencing was performed on the NovaSeq 6000 platform using SP flowcell with 2x100bp sequencing read length.

### Transcriptomic Analysis and Visualization

The raw reads were quasi-mapped against the reference transcriptomes using Salmon Quant [52]. Adaptor-trimmed reads have been uploaded to the ArrayExpress database. DESeq2 [53] was used for pairwise differential gene expression analysis between Fol-treated and untreated thalli or *in-planta* versus axenically grown fungal samples. Genes with log2FC >2 or < -2 and padj <0.05 were considered differentially up and downregulated. Custom R or Python scripts were written to visualise the results, including on chromosome level. Gene Ontology enrichment of the common DEGs across hosts genes was carried out using ShinyGO [54]. Fisher’s exact test combined with the Yekutieli (FDR) method for multiple test correction were employed for statistical testing. For Fo, ShinyGO [54] was used to perform the enrichment analysis. Here, hypergeometric tests combined with the Benjamini-Hochberg method for multiple test corrections were employed for statistical testing. In both cases, GO terms with a False Detection Rate <0.05 and p-value <0.05 were considered to be significantly enriched.

### Synteny Analysis

The proteome files of Fol and Fo47 were obtained from NCBI (GCF_000149955.1; GCF_013085055.1), and using OrthoFinder [57], single-copy orthologous genes were identified. A custom R script was written to visualise the synteny using the single copy orthogroups in a circular plot using the Circlize package. GC content and gene densities were calculated for 10-kb sliding windows across the chromosomes. In the track representing Log2FC of upregulated effectors (for the sake of visibility), the gene start and end have been multiplied by 10 to increase the effective gene length.

## ACKNOWLEDGEMENTS

We thank NGS (Next Generation Sequencing) Facility, CIFF, Electron Microscopy, IT, Greenhouse and lab-kitchen facilities at the NCBS-TIFR. We also acknowledge the help from Dr. Mugdha Sabale (Universidad de Cordoba, Spain) in assisting with the transcriptomics sampling and Dr. Christian Schudoma. (EMBL, Heidelberg, Germany) with the bioinformatic- related queries. A.R. acknowledges the support from the Ramalingaswami Fellowship, Department of Biotechnology (BT/RLF/Re-entry/72/2020) and SERB CRG grant (CRG/2022/006464), Govt. of India. This work was also supported by NCBS-TIFR intramural funding and by the Department of Atomic Energy, Government of India, under Project Identification No. RTI 4006. D.L. acknowledges the MoE-STARS/STARS-2/2023-0162 for research support.

## AUTHOR CONTRIBUTIONS

V.S. and A.R. conceptualized the work, designed the experiments. S.K., V.S., B.Z. and N.J.P. carried out the experiments and analysed the data. A.D.P. participated in the initial transcriptomic experimental design and intellectual inputs on this study. N.J.P. and D.L. contributed in generating the plant material and gave intellectual input for the *M. polymorpha* related experiments. S.K., V.S. and A.R. wrote the manuscript. All authors participated in reviewing and editing the final version of the manuscript. A.R. supervised the conducted research.

