## Supplemental Figures for "Transcriptional plasticity of fast core chromosomes governs establishment of a fungal pathogen on evolutionarily distant plant lineages"

### Supplemental Figure 1

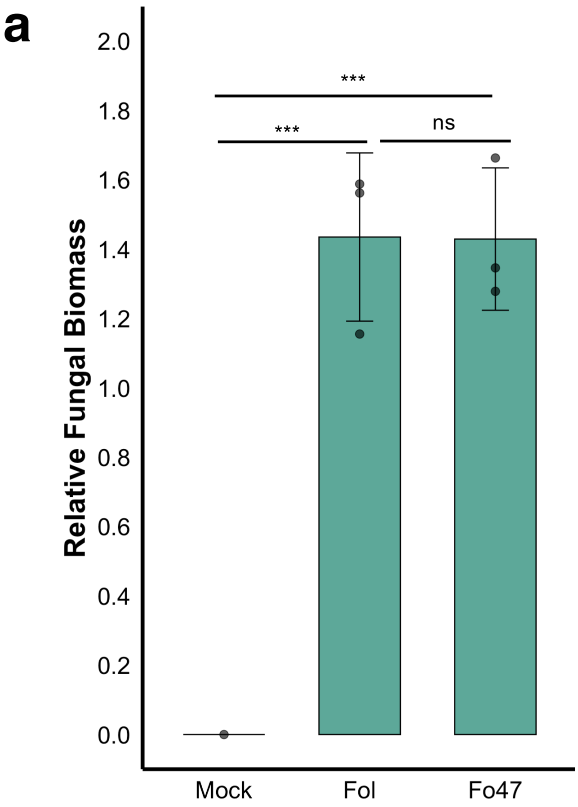


**Supplemental Figure 1: *F. oxysporum* isolates with different lifestyles infect *M. polymorpha*.** **a,** Relative fungal biomass on *M. polymorpha* Tak-1 plants 6 d after dip inoculation with the indicated Fo strains or water (mock) was measured by quantitative polymerase chain reaction (qRT-PCR) using primers specific for the Fo-*actin* gene and normalised to the *M. polymorpha* Mp*Actin* gene. Error bars indicate SD ($n = 3$). Each biological replicate represents 3 thalli. Statistical significance was calculated by one way ANOVA followed by Tukey's *post hoc* HSD test and is indicated by asterisks (***, $P < 0.001$; ns, nonsignificant, $P > 0.05$).

### Supplemental Figure 2

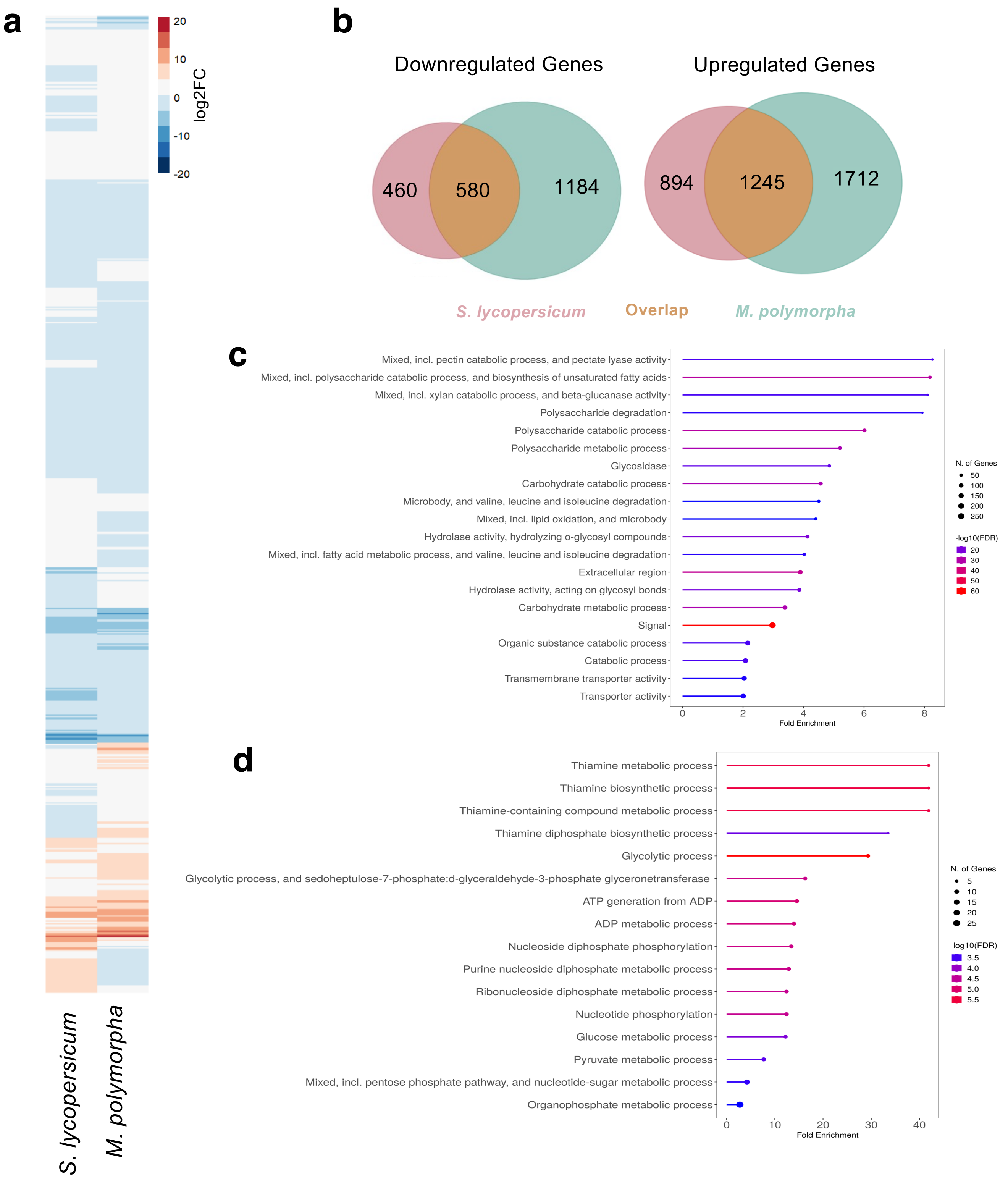


**Supplemental Figure 2: Overlap of differentially expressed genes of *F. oxysporum* during infection of evolutionarily distant plant lineages. a,** Hierarchical clustering of differentially expressed genes of *F. oxysporum* during colonization of tomato (*S. lycopersicum*) or *M. polymorpha* at 3 dpi; $p \leq 0.05$; log2 fold change [logFC] $\geq 2$. Colour scale indicates expression level. **b,** Venn diagram showing Fol DEGs during infection of tomato versus or *M. polymorpha*. **c-d,** GO analysis of genes commonly up (**c**) or down-regulated (**d**) in Fol during infection of tomato and *M. polymorpha* (FDR < 0.05).

### Supplemental Figure 3

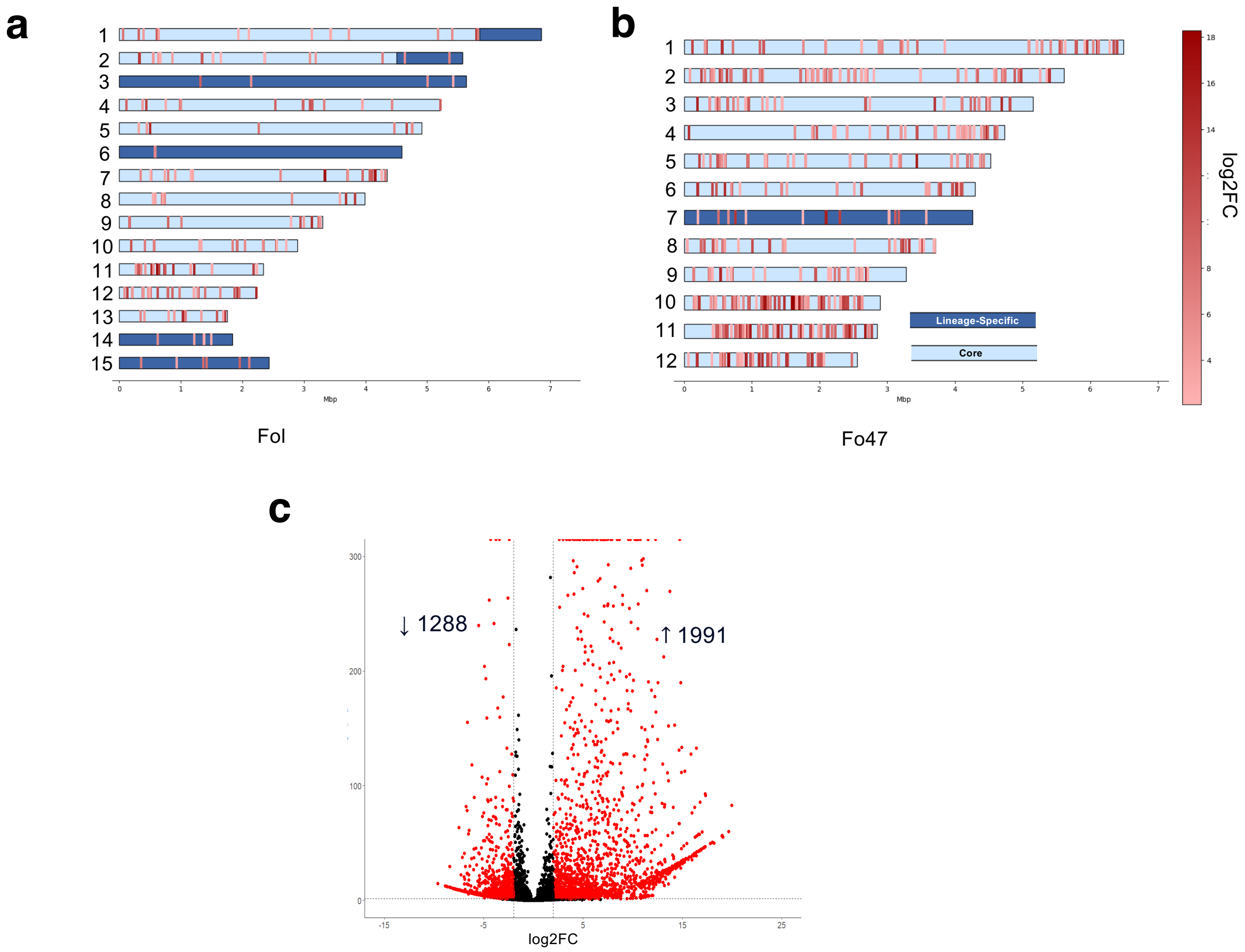


**Supplemental Figure 3: The fast core chromosomes determine compatibility of *F. oxysporum* on distant host lineages**. **a-b,** Chromosomal distribution of *in planta* upregulated genes of Fo47 **(a)** or Fol (**b**) encoding predicted secreted proteins, during infection of *M. polymorpha*. **c,** Volcano plot showing pairwise differential expression analysis of Fo47 genes suring infection of *M. polymorpha* thalli at 3 dpi compared to axenic culture. Significantly DEGs are in red.

### Supplemental Figure 4

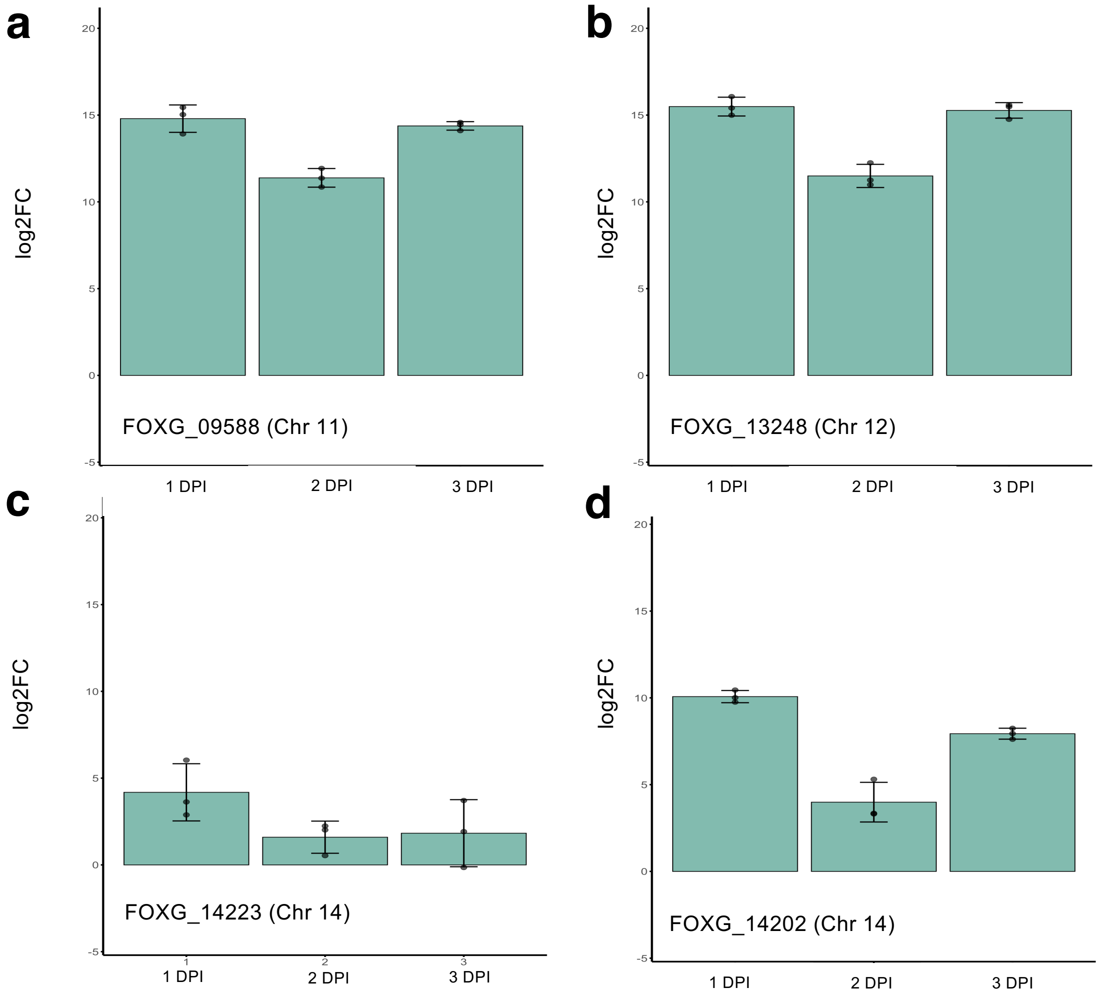


**Supplemental Figure 4: Core, but not LS effectors are upregulated in *F. oxysporum* during *M. polymorpha* infection. a-d,** Expression of effector genes in Fol wildtype during infection of *M. polymorpha* was measured at 1, 2 and 3 dpi using RT-qPCR and represented as log2 fold-change. Values shown are three biological replicates with technical replicates each, calculated as $log_2(2^{-\Delta\Delta Ct})$ using the *Foactin* gene as the internal control. Individual data points are plotted alongside mean and standard deviations.

### Supplemental Figure 5

**a**

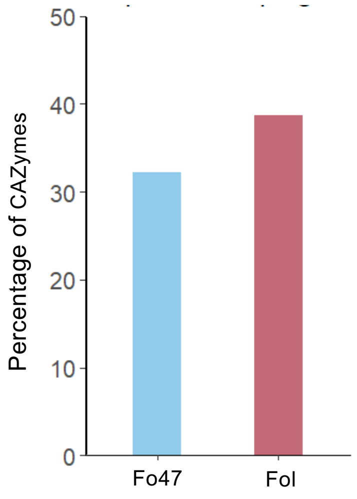


**b**

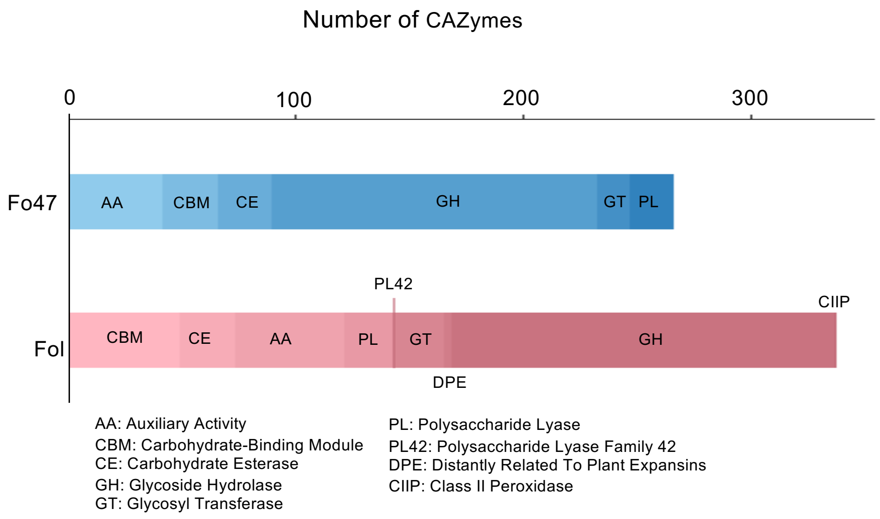


**Supplemental Figure 5: CAZYme profiles vary between Fo47 and Fol during colonization of *M. polymorpha*. a-b,** CAZyme profiles of Fo47 and Fol wildtype during infection of *M. polymorpha,* expressed as percentage of total CAZymes encoded in the respective genome **(a)** or of the families of CAZymes therein **(b).**

### Supplemental Figure 6

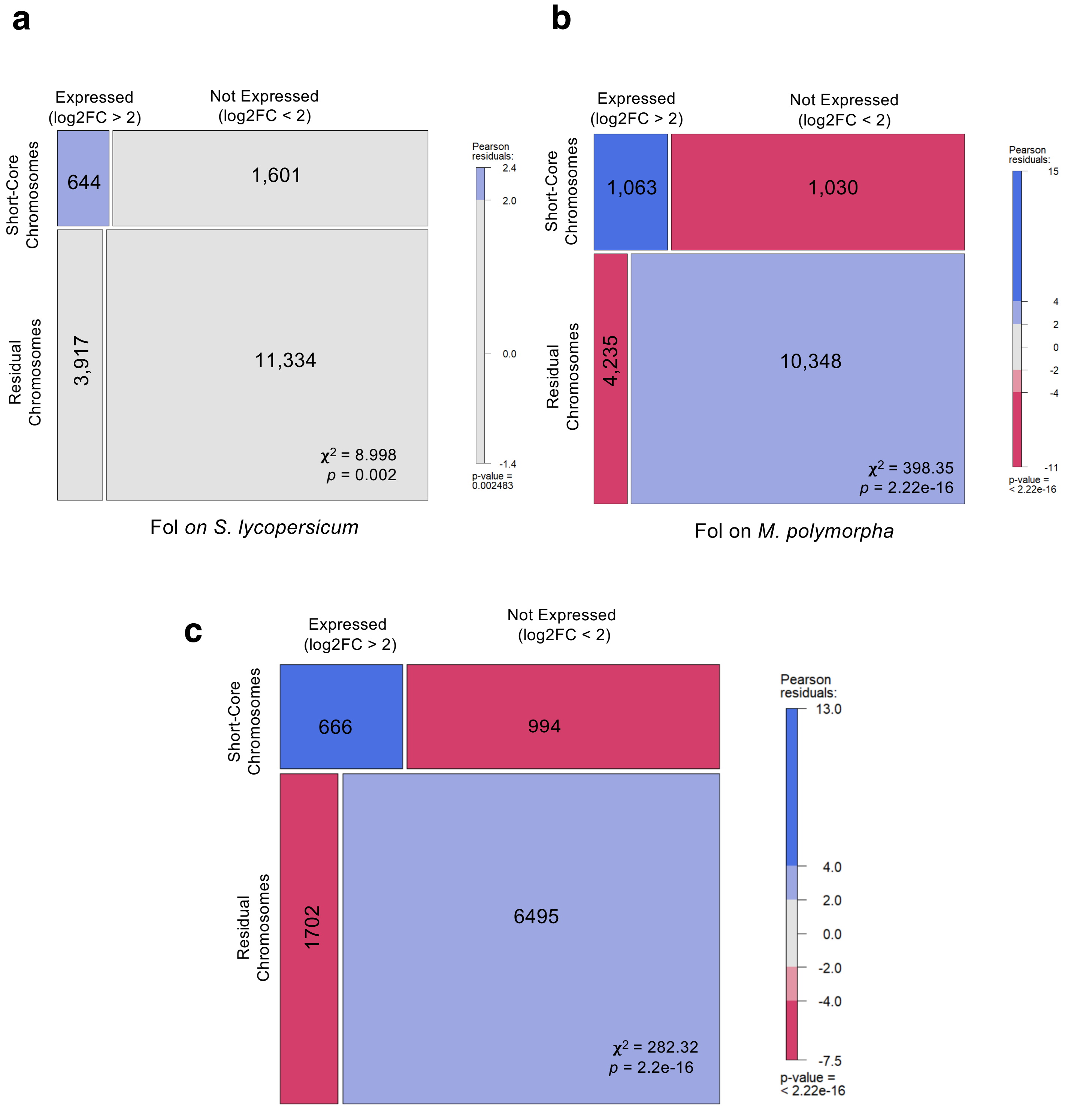


**Supplemental Figure 6: Fast core chromosomes of *F. oxysporum* are transcriptionally activated during infection of evolutionarily distant land plant lineages.**

**a-b,** Mosaic plot representing the Chi-square analysis showing enrichment of fast core genes during infection of Fol wildtype on tomato (*S. lycopersicum)* (**a**) or *M. polymorpha* (**b**). **c,** Mosaic plot representing the Chi-square analysis showing enrichment of fast core genes during infection of Fo47 on *M. polymorpha*.

### Supplemental Figure 7

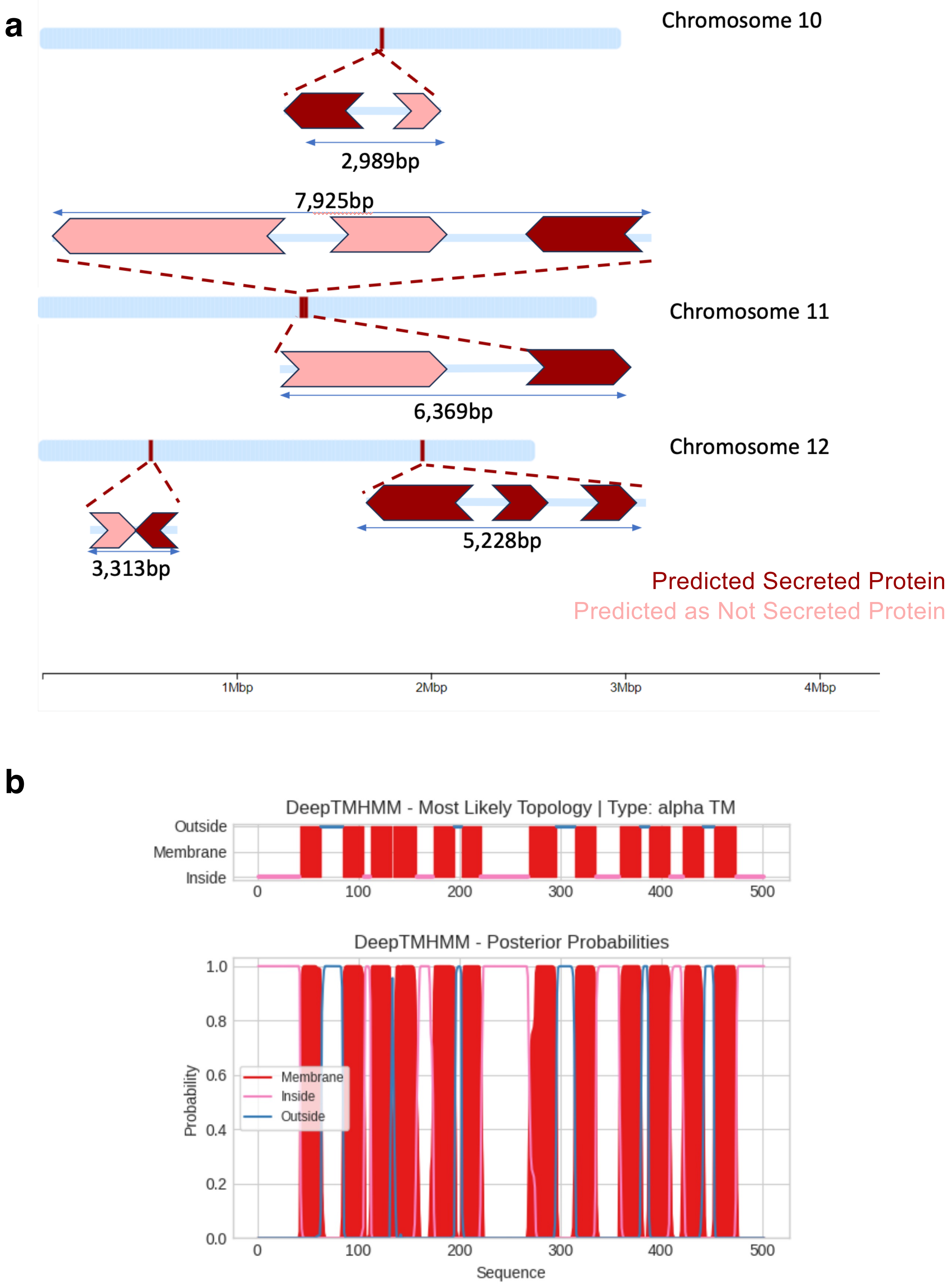


**Supplemental Figure 7: Gene clusters located on fast core chromosomes are upregulated during plant host colonization**. **a,** Schematic representation of the genomic regions on Fo47 chromosome 10, 11 and 12, comprising of five gene clusters which are transcriptionally activated during colonisation of *M. polymorpha*. Genes are depicted as arrows. Dark and light colours indicate secreted and non-secreted proteins, respectively. **b,** Predicted transmembrane domains in FOXG_16903 determined using DeepTMHMM - 1.0

### Supplemental Figure 8

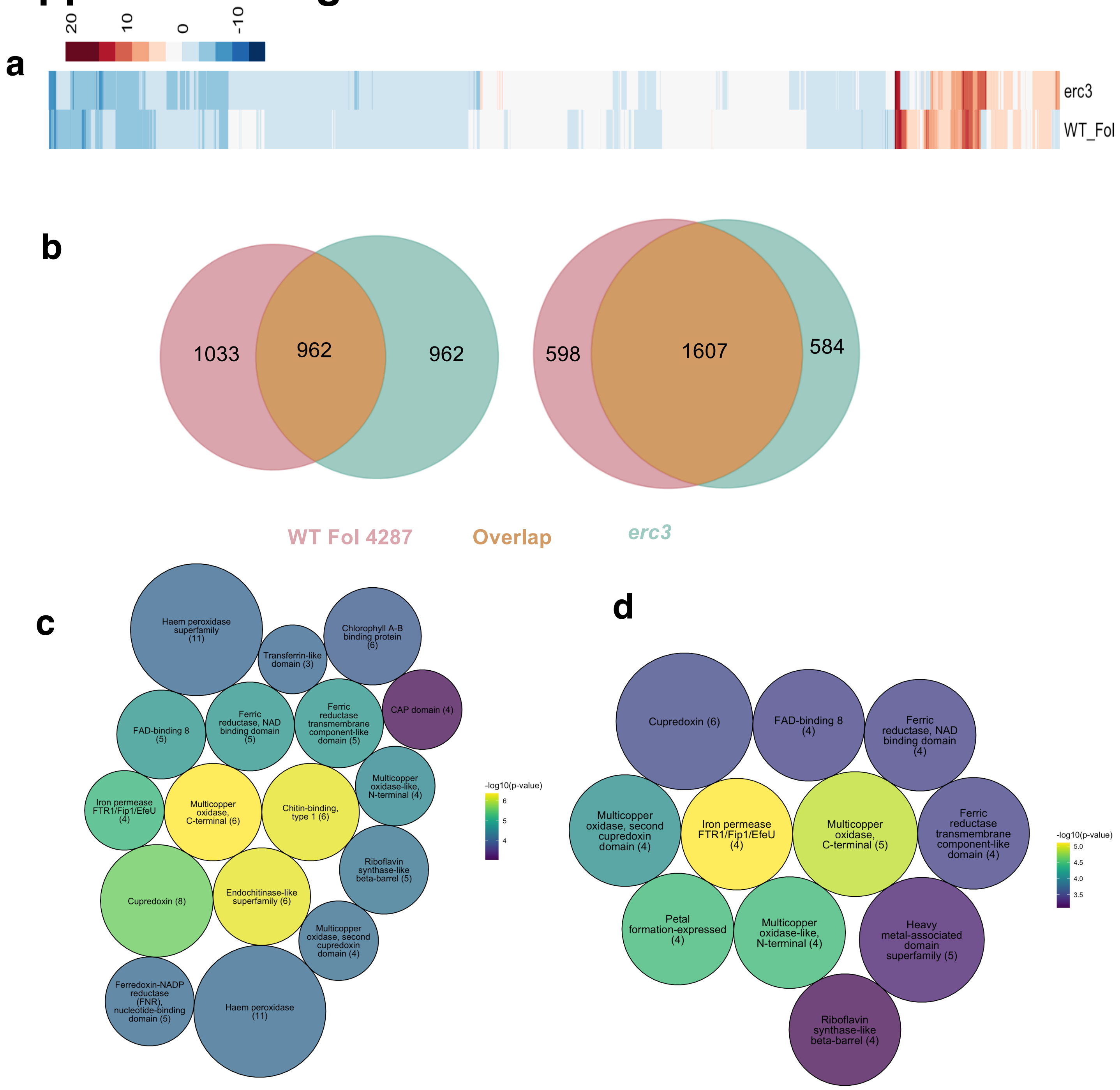


**Supplemental Figure 8: Deletion of the *F. oxysporum erc3* effector gene alters the transcriptomic landscape during *M. polymorpha* infection. a,** Hierarchical clustering of differentially expressed genes of the Fol wildtype or Δ*erc3* mutant during infection of *M. polymorpha* at 3 dpi (adjusted $p \leq 0.05$; log2 fold change [logFC] $\geq 2$). Colour scale indicates expression level. **b,** Venn diagram showing the degree of overlap between up- and downregulated genes in Fol wildtype and Δ*erc3* mutant when infecting *M. polymorpha*. **c-d,** IPR Enrichment of upregulated genes in *M. polymorpha* when infected with Fol wildtype (**a**) and *Δerc3* **(b)** using the FuncE package with an e-value cut-off of 0.001 show enrichment of chitin-binding and endochitinases only upon infection with WT Fol. The sizes correspond to the number of genes in each IPR term, and the statistical significance is represented as a -10 log of Fisher's p-value. A custom R script was written using the pack circles and the ggplot package was used to visualize the enrichment.
